# ElixirSeeker: A Machine Learning Framework Utilizing Attention-Driven Fusion of Molecular Fingerprints for the Discovery of Anti-Aging Compounds

**DOI:** 10.1101/2024.09.08.611839

**Authors:** Yan Pan, Hongxia Cai, Fang Ye, Wentao Xu, Zhihang Huang, Jingyuan Zhu, Yiwen Gong, Yutong Li, Anastasia Ngozi Ezemaduka, Shan Gao, Shunqi Liu, Guojun Li, Hao Li, Jing Yang, Junyu Ning, Bo Xian

**Author notes:** Authors contributed equally to this work (Yan Pan, Hongxia Cai). Corresponding authors: Jing Yang,; Junyu Ning,; Bo Xian.

## Abstract

Despite the growing interest in anti-aging drug development, high cost and low success rate pose a significant challenge. We present ElixirSeeker, a new machine-learning framework designed to help speed up the discovery of potential anti-aging compounds by utilizing the attention-driven fusion of molecular fingerprints. Our approach integrates molecular fingerprints generated by different algorithms and utilizes XGBoost to select optimal fingerprint lengths. Subsequently, we assign weights to the molecular fingerprints and employ Kernel Principal Component Analysis (KPCA) to reduce dimensionality, integrating different attention-driven methods. We trained the algorithm using DrugAge database. Our comprehensive analyses demonstrate that 64-bit Attention-ElixirFP maintains high predictive accuracy and F1 score while minimizing computational cost. Using ElixirSeeker to screen external compound databases, we identified a number of promising candidate anti-aging drugs. We tested top 6 hits and found that 4 of these compounds extend the lifespan of *Caenorhabditis elegans*, including Polyphyllin Ⅵ, Medrysone, Thymoquinone and Medrysone. This study illustrates that attention-driven fusion of fingerprints maximizes the learning of molecular activity features, providing a novel approach for high-throughput machine learning discovery of anti-aging molecules.

## 1 Introduction

The demand for aging retardants and prophylaxes is constantly rising as the worldwide population ages. Various pharmacological approaches have been shown to increase lifespan and attenuate age related diseases [1–8]. As anti-aging pharmacology advances, high cost and low success rate hinder drug development, emphasizing the need for identifying more promising candidate compounds. In the past, the focus has been on target-based drug discovery (TDD), which designs compounds for specific targets. The demand for more novel and promising anti-aging drug candidates would require phenotypic drug discovery (PDD), a target-agnostic approach focusing on the phenotypic impacts of treatments in relevant biological systems (Figure 1).

**Fig 1.**
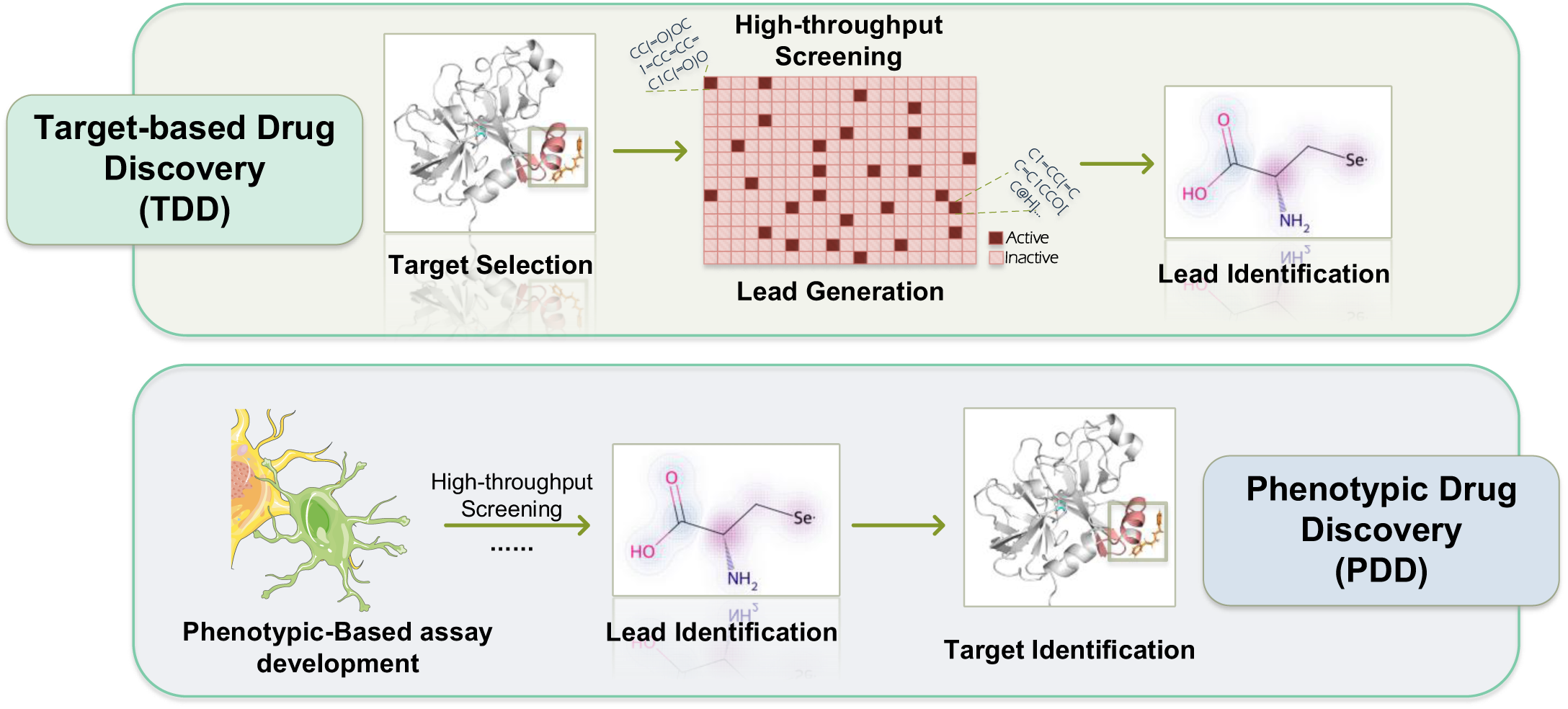
**Overview of Target-based drug discovery(TDD) and Phenotypic drug discovery(PDD)**

As a classical model organism, the nematode, *Caenorhabditis elegans* (*C. elegans*), is extensively utilized in anti-aging research. This organism is particularly valuable due to its short lifespan, simple genome, ease of manipulation, and significant genomic similarities to humans[9]. Several studies have utilized *C. elegans* to identify numerous small molecules that potentially extend lifespan and improve health. This model has facilitated the initial screening of hundreds of anti-aging compounds[10]. For example, research has demonstrated that natural products such as plant-based substances[11–12], polyphenols[13], and herbal mixtures[14] have significant potential for extending the lifespan extension of *C. elegans*. Despite the relative efficiency and high throughput of drug screening in *C. elegans*, managing the extensive number of compounds still requires substantial efforts. Thus the use of computational tools has a great potential to significantly increase the efficiency of identifying promising anti-aging compounds through pre-screening.

In the last several years, significant progress has been made in the development of solid and reliable computational chemistry and machine learning tools, reducing the time required to extract various anti-aging activities from large molecules and biology datasets[15]. AI-based techniques have been widely utilized for numerous applications, including bioactivity prediction[16], target identification[17–19], virtual drug screening[20], and drug repurposing[30]. Especially, recent advances have witnessed the utilization of various molecular fingerprints, such as MACCS, Morgan, and Topological fingerprints, in machine learning applications for analyzing untargeted metabolomics[21–23].

Molecular fingerprints are a fundamental component of drug screening methodologies, with each fingerprint based algorithm exhibiting distinct preferences. Moreover, The selection of fingerprint length, is often arbitrary; overly short fingerprints may distort critical information, while lengthy ones can introduce significant noise.[24–27]. Additionally, traditional fingerprinting approaches treat all chemical group patterns equally, without adjusting weights for specific groups based on the training set data.

To tackle these challenges, the concept of “attention-driven” fusion molecular fingerprints has been developed. This method employs weighted molecular fingerprints to enhance model precision, focusing on essential chemical groups and minimizing the influence of irrelevant data, thus boosting the efficacy of drug screening models.

Here, we provide a new machine learning framework designed to comprehensively analyze and incorporate molecular fingerprints with attention to anti-aging effect (we called “Attention-Elixir”) (Figure 2), with the goal of identifying potential small molecules with anti-aging properties. Our strategy relies heavily on the exploitation of the DrugAge database, which is a compilation of comprehensive datasets obtained from multiple investigations. We utilize the XGBoost technique to train models that determine the most effective sizes for MACCS, Morgan, and topological fingerprints. We extract from each molecular fingerprint important characteristics using XGBoost, and then use Kernel Principal Component Analysis (KPCA) to reduce the number of dimensions.

**Fig. 2.**
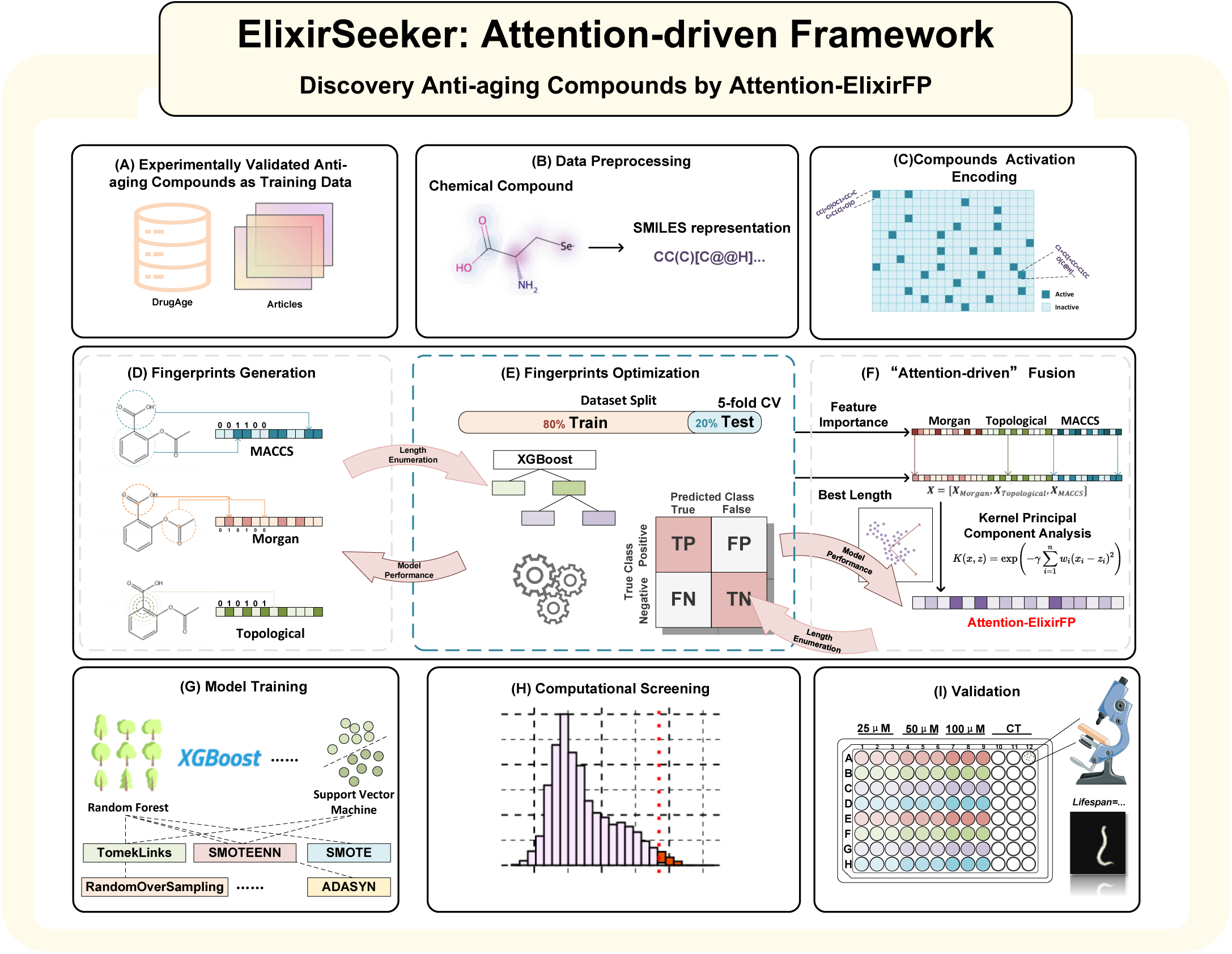
**Flowchart of the ElixirSeeker Construction.**

We applied Elixir Seeker to analyze a carefully curated database of compounds, DrugAge and identified 276 potential anti-aging drugs. We tested 6 candidates among top 0.07% compounds experimentally and found th at 4 of the candidate drugs can extend the lifespan of *C. elegans*. Committed to the principles of open science, we guarantee that all code and datasets related to this project are readily available and free to access on GitHub. Furthermore, we have included the methodologies for generating and analyzing molecular fingerprint s within the Python package, ElixirFP. This makes it easier to create customized molecular fingerprints for thorough dataset training. This tool may be easily accessed at the following URL: https://github.com/Marissapy/ElixirSeeker-ElixirFP. Its purpose is to enhance repeatability and foster additional creativity in the field.

## 2 Materials and Methods

### 2.1 Data Sources and Pre-processing

Our study leveraged data from the DrugAge database[28], which comprehensively catalogs small molecules with anti-aging properties. DrugAge aggregates information on compounds, drugs, and supplements known to extend the lifespan of various model organisms, predominantly *C.elegans*, mice, and fruit flies. This database provides detailed aging-related metrics, including average/median lifespan, maximum lifespan, strains, dosage, and gender, under standardized experimental conditions. The dataset is meticulously curated, incorporating data from diverse sources and rigorously controlled lifespan experiments. Given the substantial representation of compounds tested on *C.elegans*, our analysis primarily focused on this model organism. Additionally, we supplemented our dataset with information extracted from numerous academic publications[29–38], resulting in a dataset comprising 1695 small molecules, including 462 positive instances.

In the context of anti-aging drug screening, the problem can be abstracted into a mathematical framework, specifically as a binary classification problem. Here, the objective is to classify each molecule as either positive (having anti-aging effects) or negative (lacking anti-aging effects). Mathematically, this can be formulated using a binary label *y_i_* for each molecule *i*, where:

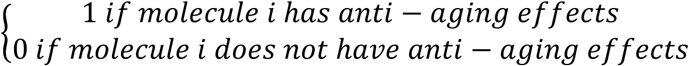

### 2.2. Feature Selection

#### 2.2.1 Generation for compounds’ fingerprints

To obtain structural information of compounds, we first used the PubchemPy tool to extract the SMILES strings of all compounds from the DrugAge database[39]. Using Python packages PubChemPy and Openbabel[40], the chemical structures of the DrugAge dataset were converted into canonical SMILES strings.

Before fusing molecular fingerprints, we conducted pre-training to determine the optimal lengths of three types of fingerprints: Morgan, Topological, and MACCS. We employed Python 3.7.10 and the following packages for generating fingerprints and training: catboost(version 1.2.5), joblib(version 1.3.2), lightgbm(version 4.3.0), numpy(version 1.22.0), pandas(version 2.0.3), rdkit(version 2023.9.5), rdkit-pypi(version 2023.9.5), requests(version 2.31.0), scipy(version 1.10.1), scikit-learn(version 1.3.2) and xgboost(version 2.0.3).

The Morgan fingerprint, also known as circular or extended connectivity fingerprint (ECFP), captures local structural features through iterative extension from each atom within a specified radius. It generates a binary fingerprint indicating the presence or absence of specific motifs, facilitating robust similarity searching and clustering. In contrast, the topological fingerprint, or path-based fingerprint, encodes molecular structures by representing topological paths or fragments as binary substructure patterns. It adeptly captures both global and local structural nuances, particularly in delineating structural similarities and pharmacophoric landscapes.

The MACCS fingerprint, derived from the Molecular ACCess System (MACCS), adopts a fixed-length representation based on predefined structural keys or pharmacophoric patterns. By identifying the presence or absence of each key, it yields a concise yet informative fingerprint.

These three types of fingerprints are calculated from different perspectives: the Morgan fingerprint captures local structural features through iterative extension, the topological fingerprint encodes molecular structures based on topological paths or fragments, and the MACCS fingerprint adopts predefined structural keys or pharmacophoric patterns for representation.

#### 2.2.2 Pre-training for Optimal Fingerprint Lengths

We utilized the XGBoost algorithm, a popular gradient boosting framework, for this task. XGBoost operates by sequentially adding decision trees to an ensemble, optimizing a differentiable loss function at each step. It uses gradient boosting to iteratively improve the model’s performance by minimizing the loss function. At each iteration, XGBoost fits a new tree to the residuals of the previous iteration’s predictions, gradually reducing the residuals and improving the model’s predictive accuracy.

One key advantage of XGBoost is its ability to provide feature importance scores, which quantify the contribution of each feature to the model’s predictive performance. Feature importance scores are calculated based on how frequently each feature is used in decision tree splits and how much each split improves the model’s performance. Features that are frequently used in important splits and lead to significant improvements in model performance are assigned higher importance scores.

Prior to feature fusion, we conducted pre-training to determine the optimal lengths of three different types of molecular fingerprints: Morgan, Topological, and MACCS. We enumerated fingerprint lengths from 16 to 1016 bits with increments of 8 (excluding MACCS), utilizing the XGBoost algorithm and employing five-fold cross-validation to obtain the optimal length for each fingerprint type. Feature importance scores were recorded for each fingerprint at its optimal length.

#### 2.2.2 Attention-Driven Fusion of Molecular Fingerprints

After obtaining feature importance scores from pre-training, we implemented an attention-driven fusion approach to integrate the fingerprints effectively.

It’s important to note that throughout the entire study, references to “ElixirFP” signify the fusion achieved using standard Principal Component Analysis (PCA) applied to the original fingerprints, while mentions of “Attention-ElixirFP” refer to the fusion accomplished through KPCA.

We used Kernel Principal Component Analysis (KPCA) with the Gaussian Radial Basis Function (RBF) kernel, leveraging the feature importance scores as weights in the KPCA.

KPCA is a dimensionality reduction technique that projects data into a higher-dimensional space defined by a kernel function. The RBF kernel used in KPCA measures the similarity between data points in the input space. The RBF kernel function is defined as:

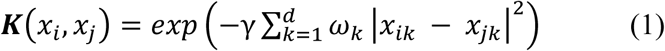

In this context, *x_ik_* and *x_jk_* represent the values of molecule i at the k-th fingerprint bit, respectively. *d* is the total length of the fingerprint, and *γ* is the bandwidth parameter of the kernel, which controls the smoothness of the Gaussian function. ω is the feature importance score of the fingerprint bit, used to amplify the differences between the fingerprint bits.

##### Algorithm 1 KPCA for Dimensionality Reduction of Molecular Fingerprints

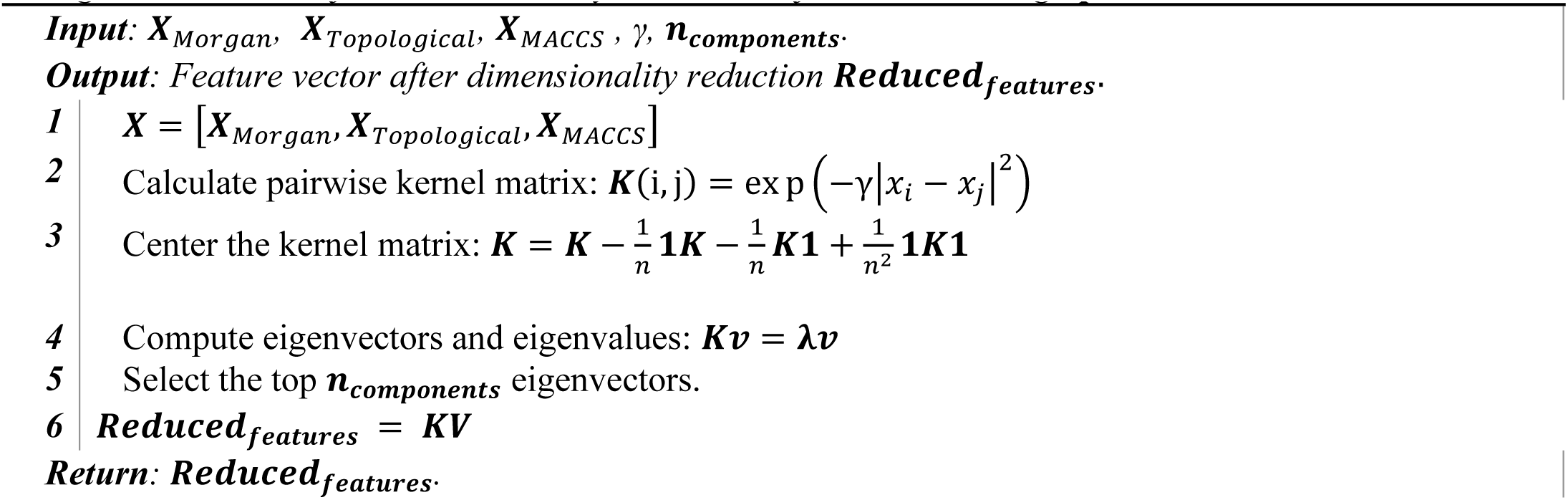

### 2.4 Model Validation

To ensure the robustness of the model, we employed a ten-fold cross-validation method. This cross-validation was conducted in scikit-learn library, with data randomly split into 80% training set and 20% test set, and five-fold cross-validation only performed within the training set. Model performance was evaluated by the AUC score, with cross-validation repeated 5 times to produce 5 AUC scores. The predictive accuracy we report is based on the average AUC values from the ten-fold cross-validation and the cumulative confusion matrix. All models were scored with three metrics of classification performance:

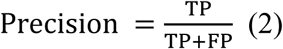

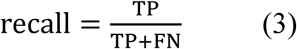

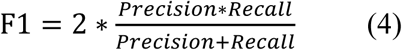

where TP represents the count of true positives, TN denotes the count of true negatives, FP is the number of false positives, and FN stands for the number of false negatives.

### 2.5 Molecular Fingerprint Stability Index (MFSI)

The Molecular Fingerprint Stability Index (MFSI) is a comprehensive metric utilized to assess the consistency of model performance across various molecular fingerprint lengths. This index is derived from the standard deviations of the model’s Accuracy, Area Under the Curve (AUC), and F1 Score under different molecular fingerprint lengths, normalized to quantify the stability of model performance. MFSI ranges between 0 and 1, where values approaching 0 indicate minimal performance discrepancies across diverse molecular fingerprint lengths, thus indicating higher stability.

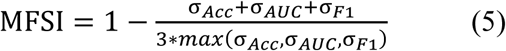

### 2.6 *C. elegans* Strains and Maintenance

In all experiments, the N2 strain was used as the wild type. Worms were maintained on NGM (Nematode Growth Medium) plates at 20°C, with the medium composed of 25 mM NaCl, 1.7% agar, 2.5 mg/mL peptone, 5 µg/mL cholesterol, 1 mM CaCl_2_, 1 mM MgSO_4_, and 50 mM KH_2_PO_4_ at pH 6.0. Escherichia coli strain OP50 was used as the food source.

### 2.7 *C.elegans* Monitoring Assay

In the process of extracting the central line from nematodes, individual specimens were imaged within a 384-well plate, allocating one nematode per well. These images were subsequently annotated using the Labelme software (version 5.4.1) within a Python (version 3.7) environment. The acquired images underwent preprocessing via the OpenCV library (version 4.9.0) in Python, which included the application of a median filter for noise reduction, enhancement of image contrast through histogram equalization, and conversion to 8-bit grayscale.

For the purpose of model development and refinement, we collected images of nematodes in liquid culture media, annotated them with Labelme, and employed data augmentation strategies to enhance the training dataset. We trained a deep learning model using this enriched dataset in PyTorch (version 2.2.0), optimizing the hyperparameters through cross-validation facilitated by the Hyperopt library.

The resulting trained U-Net model was then used to segment the nematode images, effectively isolating the central line. The segmented line was subsequently skeletonized with OpenCV techniques, refined to ensure its continuity, and purged of irrelevant branches. The finalized centerline facilitated further analysis, such as the visualization of nematode movement trajectories.

### 2.8 Lifespan Assay

Solid drug powders were dissolved in ddH_2_O or DMSO to prepare a 10 mM stock solution. The prepared drug stock was then proportionally added to the culture medium to create solutions with concentrations of 25, 50, and 100 µM. The final concentration of DMSO in the medium was maintained at 0.1% or 1%.

N2 worms were synchronized and cultured under constant 20°C conditions. Synchronized eggs were maintained at 20°C for 3 days and continued to be cultured at a constant temperature of 20°C. Worms were transferred to freshly prepared medium daily at the same time to observe their survival status. Worms confirmed as dead were removed from the medium, and the number of dead and lost worms in each culture dish was recorded. Worms lost due to crawling on the walls or handling errors, leading to non-natural death, were counted as missing.

### 2.9 Heat Shock Stress Resistance Assay

Age-synchronized 5-day-old worms were picked from NGM plates and transferred to fresh NGM plates containing drugs. After 24 hours, worms were shifted to a 35°C incubator to induce heat shock stress. Worm mortality was measured every 3 hours.

### 2.10 Oxidative Stress Resistance Assay

Age-synchronized worms, cultivated for 5 days, were selected from nematode growth medium (NGM) plates and transferred to 384-well plates containing an S-Basal solution (comprising 5.85 g/L NaCl, 1 g/L K_2_HPO_4_, 6 g/L KH_2_PO_4_, 1 mM CaCl2, and 1 mM MgSO_4_, adjusted to pH 6) supplemented with 0.1% Tween-20 and 40 mM juglone. The plates were then placed under continuous automated observation using a Zeiss microscope for a duration of 48 hours.

## 3 Results

### 3.1 Attention-driven Fusion Significantly Enhanced Model Accuracy

In our study, we conducted a comprehensive investigation of three distinct molecular fingerprinting methodologies: Morgan, Topological, and MACCS, as illustrated in Figure 2. To establish a baseline, we employed the top-performing single molecule fingerprint models. Using the Elixirseeker framework, we performed exhaustive enumeration and model training on fingerprint sizes spanning from 16 to 1016 bits. To improve the accuracy and reliability of our findings, we utilized five-fold cross-validation. Our analysis revealed that the integration of Attention-driven Fusion significantly improved model accuracy. This innovative fusion approach combined insights from diverse molecular fingerprinting methodologies.

The length of molecular fingerprints had a significant impact on training and testing precision, as well as on the AUC values and F1 scores of our models. Specifically, optimal lengths for precision analysis on the test set were determined to be 368 bits for Morgan fingerprints and 696 bits for Topological fingerprints (depicted in figure 3A-3B). To improve model performance by integrating chemical insights from diverse perspectives represented by the three fingerprints, we first subjected each fingerprint to PCA dimensionality reduction.

**Fig. 3:**
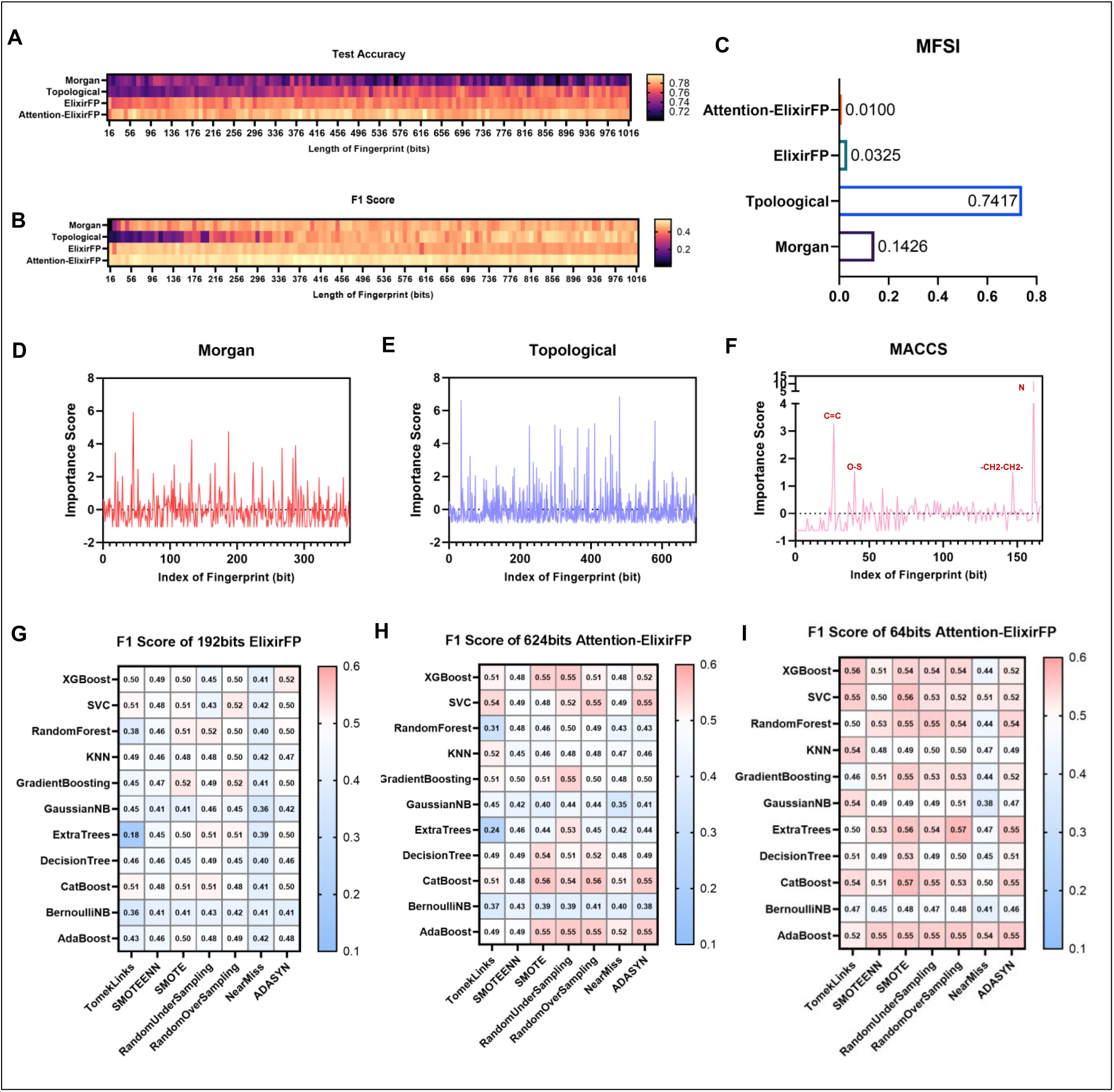
Performance Evaluation and Comparison of the ElixirSeeker Model Across Various Molecular Fingerprints and Machine Learning Techniques. (A) Heat map depicting test accuracy for Morgan, Topological, PCA, and KPCA molecular fingerprints across varying lengths from 16 to 1016 bits. (B) Heat map illustrating F1 scores for the same fingerprints across similar bit lengths, highlighting model performance consistency. (C) Bar graph showing the Molecular Fingerprint Stability Index (MFSI) for each fingerprint type, indicating the variability in model performance with different fingerprint configurations. (D-F) Feature importance scores for the optimal lengths of Morgan (D), Topological (E), and MACCS (F) fingerprints, revealing the significant bits contributing to model predictions. (G-I) Heat maps displaying F1 scores for models trained using 192-bit PCA fingerprints, and 624-bit and 64-bit KPCA fingerprints across various algorithms and sampling methods, providing a comparative analysis of the impact of algorithm and fingerprint type on the model’s predictive accuracy.

Our results indicate that the 192-bit ElixirFP generated using PCA surpassed the others in terms of accuracy (reaching 0.781). Despite higher AUC and accuracy compared to single fingerprint models, there was a slight decrease in F1 and Recall. Additionally, MFSI comparison demonstrated superior performance of the ElixirFP (figure 3C), suggesting minimal performance discrepancies across different lengths, thereby facilitating the search for optimal fingerprints.

We also introduced the concept of Attention-driven Fusion, which is a novel approach to effectively integrating molecular fingerprints. We merged these fingerprints by assigning weights to each based on their feature importance scores within the XGBoost model, and then used Kernel Principal Component Analysis (KPCA) with the Gaussian Radial Basis Function (RBF) kernel to reduce dimensionality. This attention-driven fusion approach allowed the model to focus on critical chemical properties, enhancing sensitivity to crucial structural insights and refining the ability to distinguish between active and inactive small molecules.

Incorporating previously established feature importance scores as weights resulted in a substantial enhancement in model classification performance, as illustrated in Figure 3A-3C and detailed in Table 1. KPCA exhibited superior efficacy in processing chemical data compared to standard PCA, resulting in maximal accuracy (0.824) and F1 Score (0.524) for the model using an Attention-ElixirFP.

**Tab. 1.**
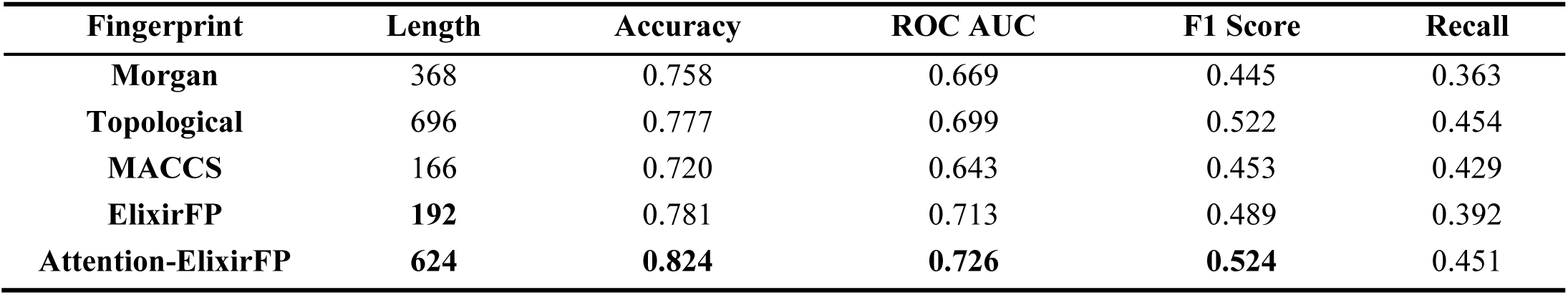
Performance Metrics of Different Molecular Fingerprints at Optimal Lengths. This table summarizes the predictive performance of various fingerprint types (Morgan, Topological, MACCS, PCA, and KPCA) at their best lengths, evaluated based on Accuracy, ROC AUC, F1 Score, and Recall.

### 3.2 Critical MACCS Fragments Driving Anti-aging Activity

In the search for powerful anti-aging compounds, a thorough analysis of key molecular fragments is essential. The MACCS (Molecular ACCess System) fingerprint is a widely used molecular descriptor that evaluates 166 substructures, and includes an additional index reserved in RDKit, increasing the total to 167 bits. It is a valuable tool for encoding the structural information of small molecules and has been extensively utilized in drug discovery and cheminformatics.

The MACCS fingerprints had particularly high significance ratings at indices 161 and 26, which were 11.41 and 3.28, respectively. These scores demonstrate the significant structural relevance these indices hold across the dataset. Index 161 indicates the presence of a nitrogen atom, as depicted in figure 2F, while index 26 indicates a carbon atom in a cyclic structure with certain double bonds and connection patterns. These specific chemical structures play a pivotal role in the predictive modeling of the anti-aging activity of small molecules, highlighting their crucial contribution to the model’s accuracy.

Further investigation revealed that two of the top five ranked fragments notably include nitrogen atoms, as illustrated in Table 2. This emphasizes the pivotal role of nitrogen in the formation of bioactive molecules, as well as its significance in distinguishing between active and inactive small molecules. The chemical versatility of nitrogen which allows it to form a wide range of bonds such as amines, amides, and imines, makes it an essential component of many biologically active compounds.

**Tab. 2.** Key structural motifs identified using MACCS (Molecular ACCess System) indices. . This table presents the MACCS indices with the highest importance scores, along with their corresponding SMARTS notations and descriptions of the molecular structures they represent. The indices illustrate crucial chemical features that significantly contribute to the anti-aging properties of compounds.

| MACCS Index | Importance Score | SMARTS | Description |
| --- | --- | --- | --- |
| 161 | 11.411 | [#7] | Nitrogen atom |
| 26 | 3.238 | [#6]=;@[#6](@*)@* | Structure of a carbon-carbon double bond (C=C) between two carbon atoms |
| 40 | 1.526 | [#16]-[#8] | Sulfur-oxygen single bond |
| 147 | 1.495 | \$(*~[CH2]~[CH2]~*),\$([R]1@[CH2;R]@[CH2;R]1) | A methylene bridge (-CH2-CH2-) linked to any other atom |
| 25 | 0.964 | [#7]~[#6](~[#7])~[#7] | Structure of a carbon atom directly connected to three nitrogen atoms |

### 3.3 64-bits Reduced Dimensionality Fingerprints Maintain High Performance

In experiments involving dimensionality reduction of various molecular fingerprints by the Elixirseeker framework, we observed that a 64-bit Attention-ElixirFP achieved a test accuracy of 79.4%, compared to 72.6% and 73.1% for equivalent-length Morgan and Topological fingerprints, respectively (see Table 3). While significantly reducing dimensionality, we successfully enhanced accuracy. Notably, the performance of the 64-bit Attention-ElixirFP was only 3% lower than our identified optimal molecular fingerprint (624-bit Attention-ElixirFP). Given the trade-off between accuracy and model complexity, this minor discrepancy is acceptable. The 64-bit Attention-ElixirFP reduced by KPCA not only substantially decreased feature dimensions and computational costs, but also retained high-precision representation capacity for anti-aging small molecules.

**Tab. 3.** Performance of Different Molecular Fingerprints at a Length of 64 Bits. This table showcases the predictive performance metrics for four types of fingerprints (Morgan, Topologica, ElixirFP, and Attention-ElixirFP) reduced to a uniform length of 64 bits. The Attention-ElixirFP 64-bit fingerprint demonstrates superior performance across all metrics, highlighting its efficacy in capturing essential molecular features necessary for anti-aging compound identification.

| Fingerprint | Length | Accuracy | ROC AUC | F1 Score | Recall |
| --- | --- | --- | --- | --- | --- |
| Morgan | 64 | 0.726 | 0.651 | 0.435 | 0.392 |
| Topological | 64 | 0.731 | 0.546 | 0.157 | 0.095 |
| ElixirFP | 64 | 0.769 | 0.696 | 0.487 | 0.412 |
| Attention-ElixirFP | 64 | **0.794** | **0.707** | **0.534** | **0.440** |

Based on the disparity in the number of positive and negative class samples in our training set, we expanded our research to test ensemble fingerprints. We chose the F1 score as the primary metric for training models rather than relying solely on accuracy in order to provide a more balanced evaluation of model performance. This approach is particularly critical in circumstances with skewed class distributions, when accuracy alone may not adequately reflect the true effectiveness of a model.

Furthermore, we employed 192-bit ElixirFP fingerprints, as well as 64-bit and 624-bit Attention-ElixirFP fingerprints, to evaluate a combination of twelve common machine learning models and seven sampling techniques, as illustrated in Figure 3G-3I. The findings from these tests indicated that the 64-bit Attention-ElixirFP fingerprints produced the best overall performance. Notably, when combined with the optimal ‘ExtraTrees + RandomOverSampling’ model, the 64-bit Attention-ElixirFP fingerprint recorded the highest F1 score of 0.572 (Table 4). This score represents a 5% increase in the F1 score and a 7.7% increase in recall over the previous best-performing single fingerprint model, which utilized a 696-bit Topological fingerprint.

**Tab. 4.** TOP5 best model performance of ElixirFP and Attention-ElixirFP Fingerprints with Different Machine Learning Models and Sampling Techniques at 64 and 624 Bits. This table details the results of various combinations of machine learning models and sampling methods using Attention-ElixirFP fingerprints at both 64 and 624 bits. It evaluates performance across Accuracy, ROC AUC, F1 Score, and Recall. The ExtraTrees model combined with RandomOverSampling and the 64-bit Attention-ElixirFP exhibits the highest F1 Score and substantial Recall, indicating its optimal balance for handling imbalanced datasets and enhancing model precision and recall. Note: The 192-bit ElixirFP is not included in the top 5 rankings, hence it is not listed here.

| Fingerprint | Model | Length | F1 Score | ROC AUC | Accuracy | Recall |
| --- | --- | --- | --- | --- | --- | --- |
| Attention-ElixirFP | ExtraTrees+RandomOverSampling | 64 | 0.572 | 0.742 | 0.786 | 0.531 |
| Attention-ElixirFP | CatBoost+SMOTE | 64 | 0.569 | 0.718 | 0.769 | 0.560 |
| Attention-ElixirFP | SVC+SMOTE | 64 | 0.561 | 0.722 | 0.773 | 0.532 |
| Attention-ElixirFP | CatBoost+SMOTE | 624 | 0.559 | 0.711 | 0.770 | 0.545 |
| Attention-ElixirFP | ExtraTrees+SMOTE | 64 | 0.559 | 0.728 | 0.779 | 0.520 |

### 3.4 Key Molecular Signals Determining Anti-Aging Properties

To explain why the PCA method enhances the model, as described earlier, we aim to find the important signals driving anti-aging activity of each molecular fingerprints. As shown in Figure 4A, Fingerprint information from lengths 1057 to 1100 is insignificant, indicating redundancy and noise. In contrast, fragments from 1101 - 1231, indicated by MACCS fingerprints, exhibit higher loading values, emphasizing their crucial role in indicating biological activity.

**Fig. 4:**
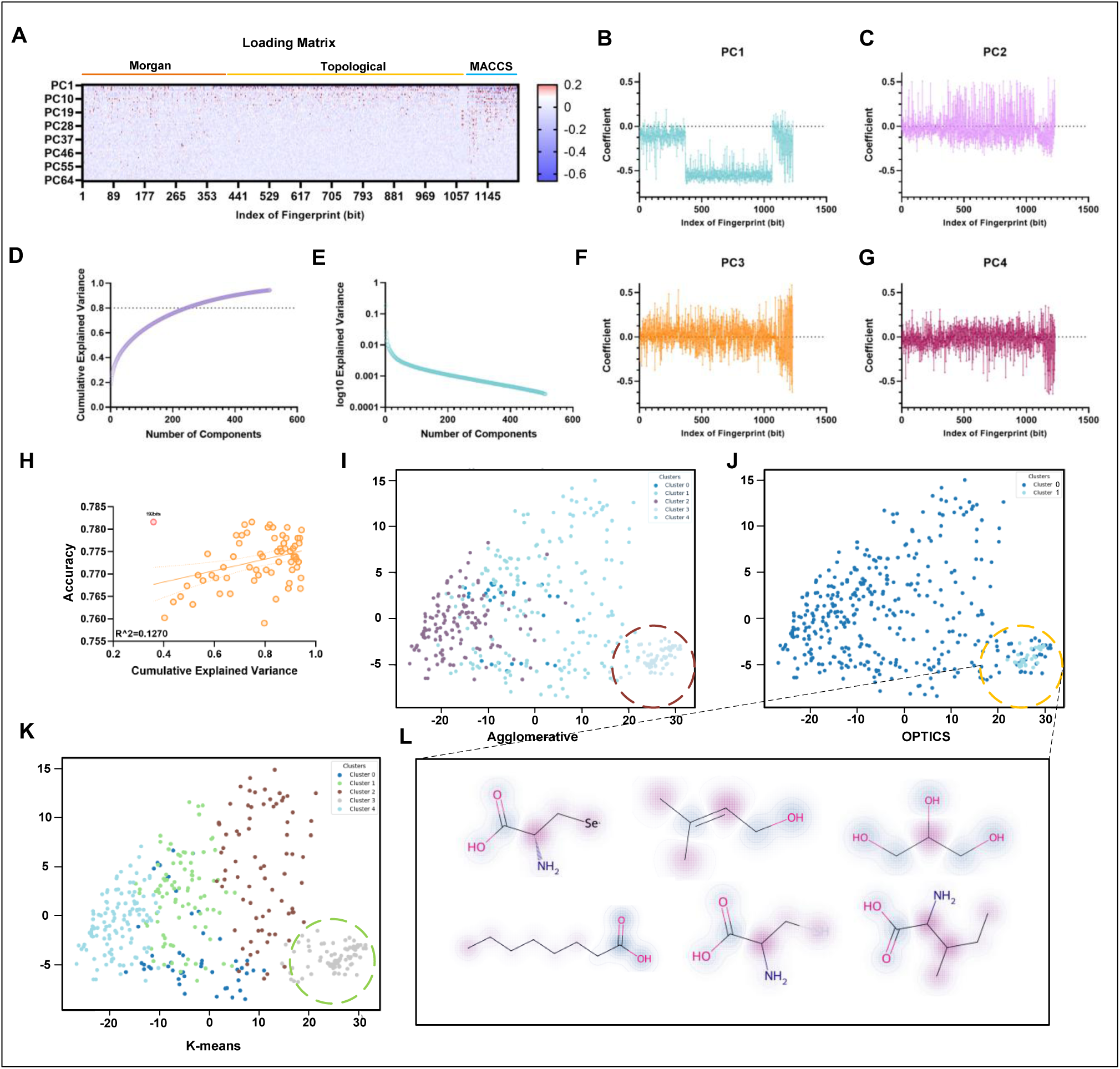
Detailed Analysis of PCA Components and Clustering of Molecular Fingerprints in Anti-Aging Compound Discovery. (A) Loading matrix for the 64 components across Morgan, Topological, and MACCS fingerprints, highlighting the significant bits contributing to each component. (B-G) Coefficient plots for principal components PC1 (B), PC2 (C), PC3 (D), and PC4 (E), detailing the contribution of individual fingerprint bits to each principal component across the entire spectrum. (H) Scatter plot depicting the relationship between cumulative explained variance and model accuracy (I-K) KPCA-based clustering results, showing distinct clusters of molecular compounds colored by class: (I) Agglomerative clustering, (J) OPTICS clustering, and (K) K-means clustering. Each plot identifies nuanced grouping within the chemical space, with highlighted clusters indicating unique or significant features. (L) Representative molecular structures from key clusters identified in the OPTICS method.

As shown in Figure 4A-4C and 4F-4G, the first principal component (PC1) has large and consistent negative coefficients, indicating common trends among features. Despite their diverse contributions, these features collectively highlight shared chemical properties. The coefficients distribution of the second to fourth principal components is more uniform, with relatively smaller values, indicating balanced feature contributions and slight structural changes.

Figure 3D-3E demonstrate that the first 64 principal components capture approximately 50% of the cumulative explained variation while maintaining good prediction accuracy. This indicates that these components effectively capture important anti-aging activity signals. Furthermore, we find essentially no correlation between cumulative explained variance and accuracy (Figure 4H). Despite having a relatively lower explained variance, ElixirFP generated by PCA significantly enhances model accuracy, suggesting that the anti-aging efficacy of small molecules is driven by key signals that are efficiently captured by these principal components. This emphasizes the importance of using weighted KPCA for dimensionality reduction of molecular fingerprints, as it can reduce redundant features and noise, allowing the model to focus on important variables. Such an approach is valuable for screening bioactive small molecules in drug discovery, highlighting that key signals are often determined by a small subset of features.

### 3.5 Clustering Analysis of anti-aging compounds

As we delve deeper into understanding the landscape of anti-aging compounds, clustering analysis emerges as a powerful tool for unraveling intricate relationships among molecules. In this study, we characterized a series of anti-aging small molecules in the train set using 64-bit Attention-ElixirFP, to reveal subtle similarities and significant differences among these molecules.

We employed two distinctly different clustering algorithms: OPTICS, which represents unsupervised learning, and K-means and Agglomerative Clustering, which are commonly used in supervised learning. Consistent with previous studies, these methods show that anti-aging small molecules are difficult to categorize based on common characteristics. As shown in figures 4I-4K, the OPTICS algorithm distinguished two significant groups of small molecules, but K-means and Agglomerative Clustering categorized the molecules into five classes on a finer scale. Different clustering methods for classifying anti-aging small molecules allow us to comprehend the interactions between molecules from different perspectives and infer possible biological properties based on their distribution in chemical space.

It is worth noting that, regardless of the clustering method used, molecules on the right side of the diagram are consistently classified as a separate group, suggesting a fundamental similarity in their chemical structures or biological functions. The tendency of these molecules to cluster in specific areas may indicate that they possess certain key biological activities or molecular properties.

In the aforementioned category (i.e., classified as Class 1 by OPTICS clustering), we observe that most molecules with anti-aging properties contain the carboxyl group (-COOH), which prevalence may be related to its versatile biological functions in the body, as shown in table 5. Carboxylic acids, as a critical component of cellular metabolism, play an important role in multiple key biochemical pathways, including but not limited to functioning as metabolic intermediates, maintaining intracellular pH equilibrium, and participating in cell signaling. These characteristics make small molecules containing carboxyl groups ideal candidates for exploring anti-aging mechanisms. Therefore, the presence of the carboxyl group may be linked to their potential anti-aging effect, providing a valuable chemical foundation for the development of future anti-aging drugs.

**Tab. 5.**
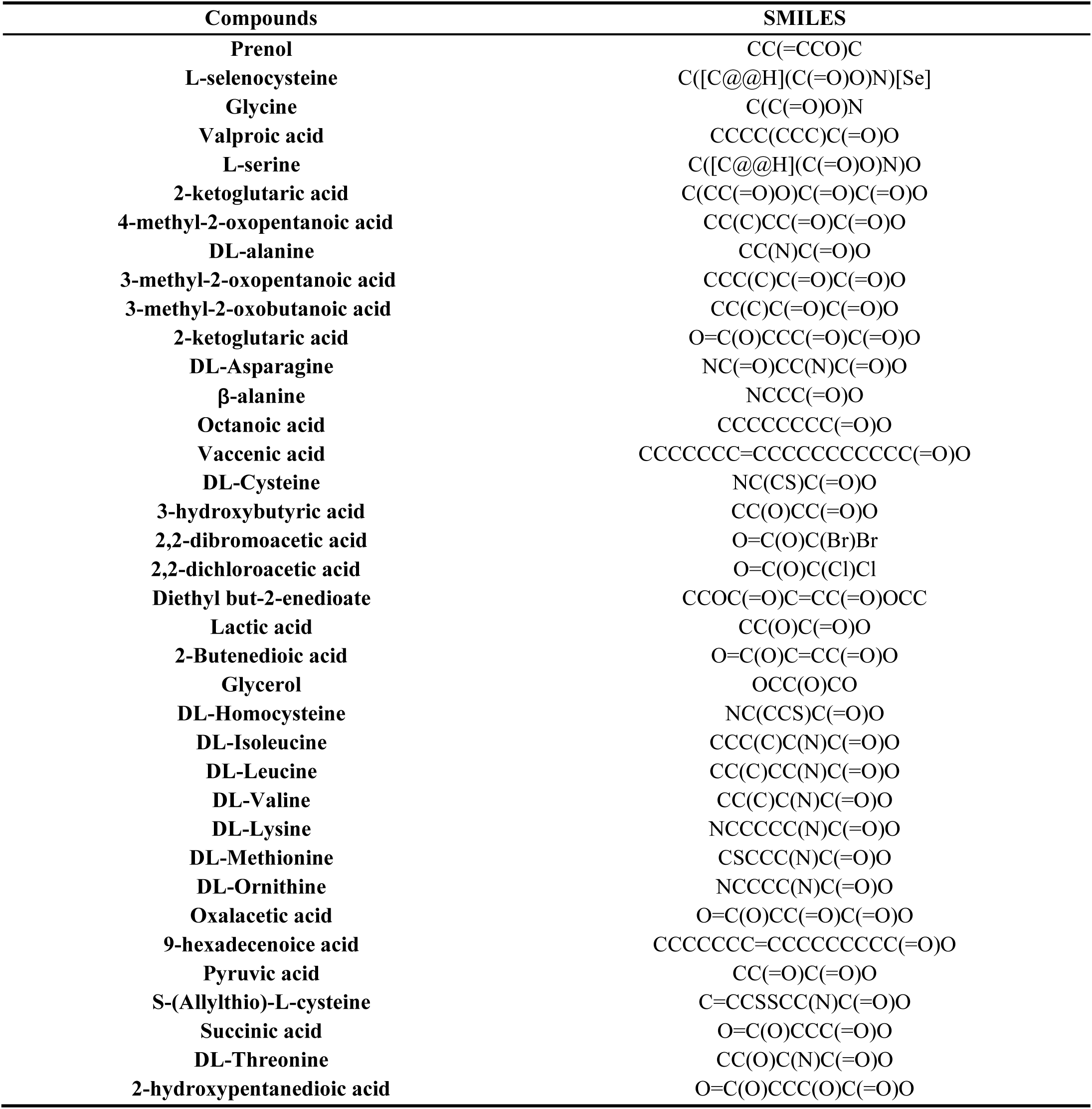
Compounds from key clusters identified in the OPTICS method.

**Tab. 6.** Effects of Thymoquinone (TQ) on Lifespan and Heat Shock Resistance in *C. elegans*. Mean lifespan and heat shock resistance (in hours) for *C. elegans* treated with varying concentrations of Thymoquinone (25μM, 50μM, and 100μM) relative to the control (ddH2O). Data are expressed as mean ± standard error of the mean (SEM). Statistical significance was assessed using the Log-rank test for survival data, with p-values indicated. Hazard ratios (HR) and 95% confidence intervals (CI) of the ratio are also provided. Asterisks (*) denote levels of statistical significance compared to the control group: *p < 0.05, **p < 0.01, ***p < 0.001, ****p < 0.0001.

| Treatments | Lifespan |  |  |  |
| --- | --- | --- | --- | --- |
|  | Mean±SEM | Log rank p | HR | 95% CI of ratio |
| ddH<sub>2</sub>O | 12.54±0.44 | - | - | - |
| 25µM | 14.21±0.34 | 0.0051** | 0.797 | 0.599 ~ 1.060 |
| 50µM | 14.34±0.41 | 0.0078** | 0.752 | 0.566 ~ 0.998 |
| 100µM | 14.21±0.34 | 0.1261 | 0.910 | 0.677 ~ 1.224 |

**Tab. 6 Effects of Thymoquinone (TQ) on Lifespan and Heat Shock Resistance in *C.**
| Treatments | Heat Shock |  |  |  |
| --- | --- | --- | --- | --- |
|  | Mean±SEM | Log rank p | HR | 95% CI of ratio |
| ddH<sub>2</sub>O | 36.06±2.39 | - | - | - |
| 25µM | 43.05±3.10 | 0.0317* | 0.687 | 0.456 ~ 1.034 |
| 50µM | 49.37±2.45 | <0.0001**** | 0.537 | 0.359 ~ 0.804 |
| 100µM | 44.09±2.87 | 0.0197* | 0.669 | 0.445 ~ 1.004 |

### 3.6 Model Application for Compound Profiling and Screening

After optimizing and integrating molecular fingerprints, we expanded the application of our model to an external compound database to assess its generalizability and effectiveness outside the initial training set. Using the 64-bit Attention-ElixirFP developed in this study, our screening process yielded notable results, particularly with the ‘ExtraTrees + RandomOverSampling’ ensemble method as the final ElixirSeeker Model. Notably, we identified several potential compounds with high prediction confidence.

Our screening process included three compound libraries: the FDA approved library (TargetMol, USA, L1010), the Sellect Bioactive compound library (Selleckchem, Houston, TX, USA, L1700), and a Traditional Chinese Medicine compound library (Chengdu Biopurify Phytochemicals Ltd.), which contained 3151, 9109, and 1988 small molecules, respectively. Using our model for prediction, we obtained a maximum score of 0.716. We then selected the top 2% of small molecules with a threshold score of 0.57(Figure 5A, S1, S2, S3). Most compounds were assigned prediction scores indicating a low likelihood of being senolytic-like. Further analysis of the compounds over this threshold yielded intriguing findings. First, none of the top 2% of small molecules occurred in the training set, indicating their novelty. Additionally, some molecules had previously been validated as positive compounds, such as Triptolide[41] and 20-Deoxyingenol[42].

**Fig. 5:**
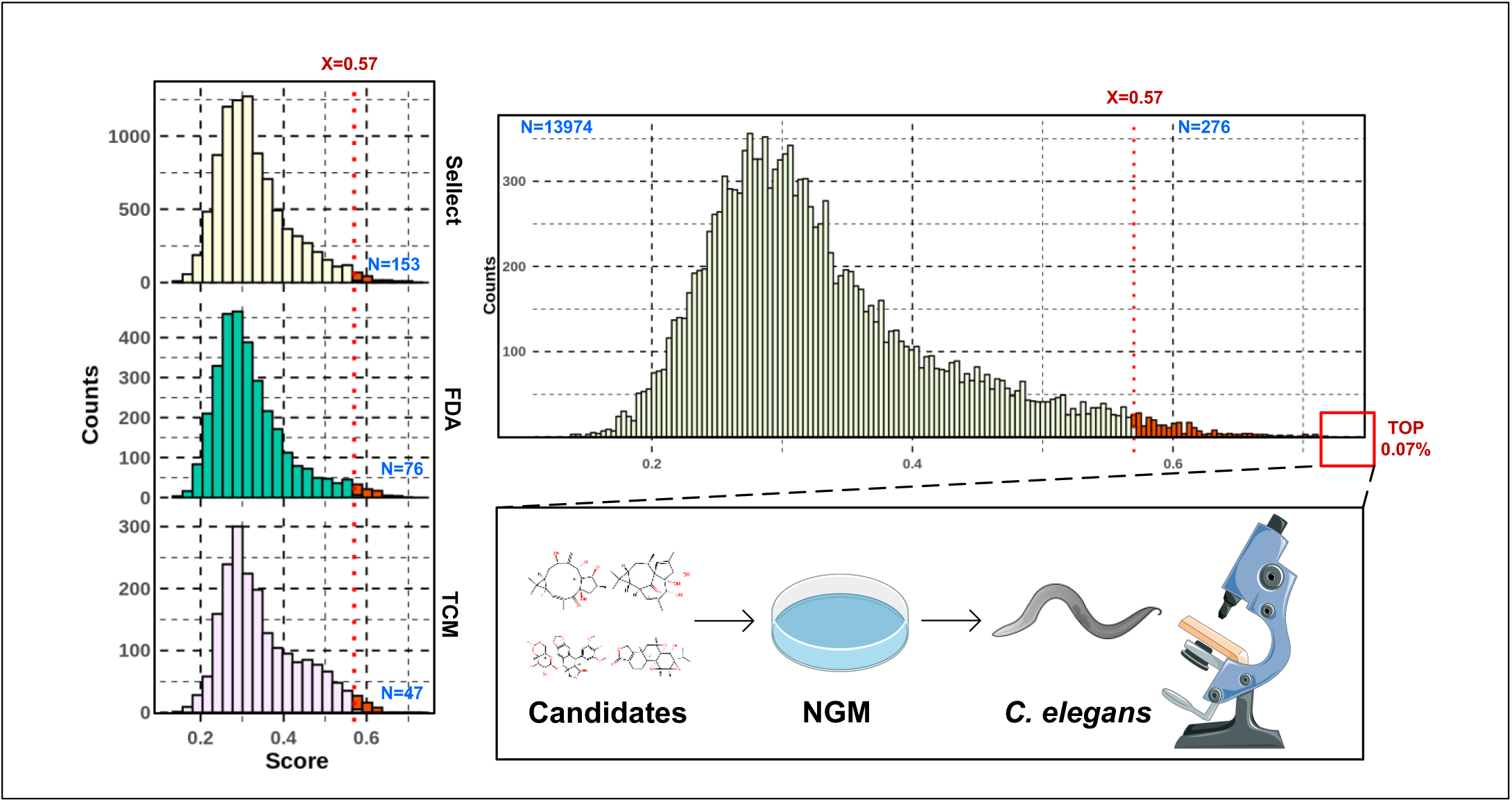
External database screening and validation results. Density distribution of scores from screening the Sellect, FDA, and TCM databases using the Elixirseeker model.

The composition of small molecules with a score greater than 0.57 in the Traditional Chinese Medicine compound database is particularly interesting. Notably, 89.36% of these compounds belong to the saponin class, which is known for its anti-aging properties. Specifically, 40% of these saponins are Ginsenosides, including Ginsenoside F1, highlighting the relevance of Ginseng in anti-aging research.

Moreover, in the Traditional Chinese Medicine database, Thymoquinone emerged with a notably high score of 0.696, surpassing the second-ranked Ginsenoside-Rh4 (0.637). This prompted our interest in investigating whether Thymoquinone could extend the lifespan of *C.elegans*, hence revealing its potential as an anti-aging compound. Although reports have suggested the activity of Thymoquinone, no study has demonstrated that it extends lifespan in any model organism. As a result, we are particularly intrigued by the potential of Thymoquinone in this area.

Then, we conducted in-depth biological activity tests on a selection of top-performing small molecules identified from the three libraries, which had not been previously reported to have anti-aging activities. These molecules included Praeruptorin C, Polyphyllin VI, α-Hederin, Medrysone, Thymoquinone, and 7β-Hydroxylathyrol. These compounds were chosen based on their high prediction scores and their absence in literature concerning direct anti-aging or senolytic activities, making them prime candidates for new research.

### 3.7 Selected Compounds Enhance *C. elegans* Survival Under Heat Stress

Firstly, we conducted heat shock stress experiments on *C. elegans* at 37 °C. Heat shock experiments serve as supportive evidence for lifespan assays, enabling rapid conclusions about their relationship with anti-aging mechanisms.

All exhibited trends toward extending the lifespan of *C. elegans* under heat shock stress conditions. However, since the concentration of each small molecule was uniform across experimental groups, some compounds were either too high or too low in concentration to exhibit this trend. Nonetheless, this still demonstrates their molecular activity and potential anti-aging properties, as illustrated in Figure 6.

**Fig. 6:**
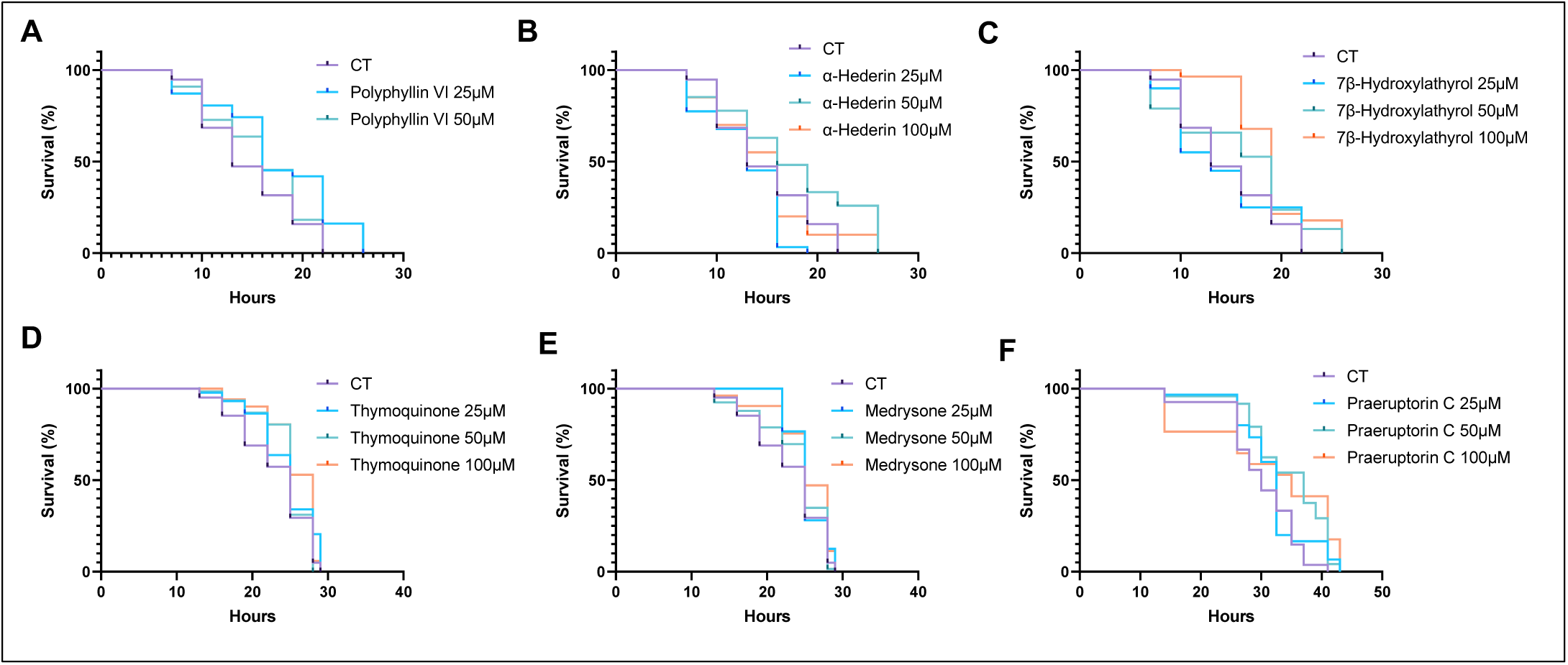
**Survival curves under heat shock stress after five days of treatment with candidate compounds.**

We are finding that among the compounds tested, especially 7β-Hydroxylathyrol at a concentration of 100 μM, significantly extends the lifespan of the worms by 46% under heat shock stress conditions (p=0.003). It suggests that the compound improves the organism’s resilience to acute stressors, which is a key feature of potential anti-aging substances.

### 3.8 Candidate Compounds Extends the Lifespan in *C. elegans*

Further, we conducted lifespan assays on *C. elegans* using six candidate compounds. Our findings reveal that four of these small molecules significantly extend lifespan, specifically Polyphyllin VI (Figure 7A), Thymoquinone (Figure 7D), Medrysone (Figure 7E), and Praeruptorin C(Figure 7F). Notably, 7β-Hydroxylathyrol showed the greatest increase in maximum lifespan (Figure 7C). Polyphyllin VI, in particular, demonstrated the most pronounced effect on lifespan extension, increasing longevity by 20% at a concentration of 25μM (p<0.0001). Polyphyllin VI is an active saponin derived from the traditional medicinal plant Paris polyphylla.

**Fig. 7:**
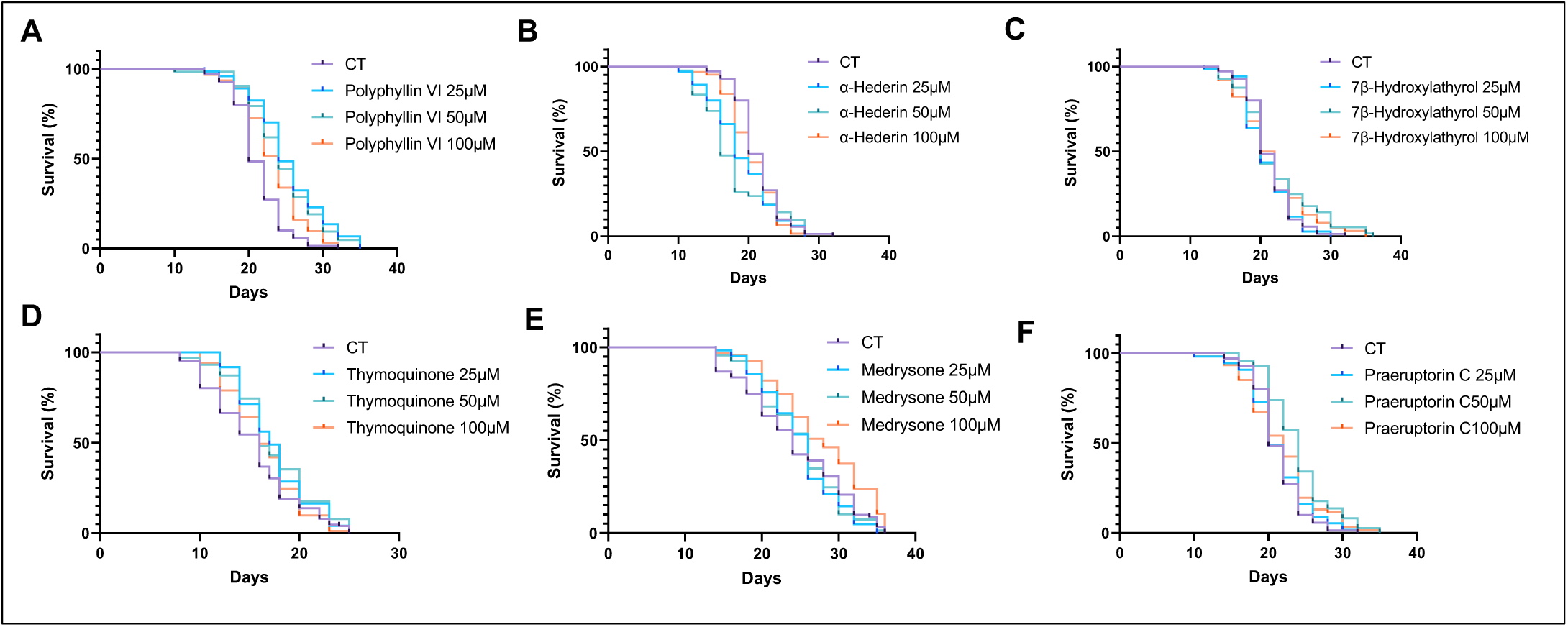
**Lifespan curves for nematodes treated with candidate compounds.**

Similarly, Thymoquinone, a bioactive compound extracted from Nigella sativa—commonly known as black cumin[43]—also exhibited significant lifespan extension effects at a 25μM concentration. Further discussion on these findings will be addressed in the Discussion section of the paper.

## 4 Discussion

### 4.1 Importance of Molecular Fingerprint Optimization and ‘Attention-driven’ Fusion

The careful refinement and combination of molecular fingerprints are crucial in modern drug development to improve the effectiveness and accuracy in identifying potential candidate drugs. Molecular fingerprinting techniques condense intricate chemical structure information into readily understandable numerical data, facilitating the efficient processing and examination of extensive compound libraries by machine learning algorithms.

However, as previous studies have shown, capturing the characteristics of small molecules with anti-aging effects, which often exhibit subtle similarities, has proven difficult. Traditional molecular fingerprinting methods usually encounter challenges such as high dimensionality and information redundancy, which can potentially increase computational demands while decreasing model prediction accuracy and generalizability. In recent years, numerous new molecular fingerprints have been developed specifically for deep learning and other advanced methods [44–45], however they frequently require substantial computational resources.

This work enhances the DrugAge database by incorporating previous research and introducing other novel aspects. To effectively address the issues posed by the complex characteristics of small compounds with anti-aging properties, the model achieves a wider understanding of the diversity in chemical space by incorporating many types of molecular fingerprints, such as Morgan, topological, and MACCS. This not only captures more intricate chemical structure changes but also more precisely portrays the intricate interactions among molecules.

Second, the concept of “attention-driven fusion” in ElixirSeeker is proposed. Incorporating feature weights derived from XGBoost’s “feature importance” scores into KPCA creates a weighted analysis technique. KPCA, particularly with the Gaussian Radial Basis Function (RBF) kernel, broadens the possibilities of traditional PCA. This kernel excels in transforming the original nonlinear feature space into a high-dimensional space in which principal components are linearly separable.

This transformation is crucial for modeling the inherent complex and often nonlinear molecular interactions in pharmacological studies. In this framework, more significant features exert greater influence on the model, enabling KPCA to prioritize data elements that most effectively predict biological activity.

Furthermore, we introduced the concept of “MFSI” (Molecular Fingerprint Stability Index). Because determining the optimal length of screening molecular fingerprints requires substantial computational power, the Molecular Fingerprint Stability Index (MFSI) is proposed as a means of evaluating the stability of molecular fingerprints. MFSI serves as a tool in mitigating the computational burden associated with determining optimal fingerprint lengths.

Fourth, through an elucidation of the principles underlying Principal Component Analysis (PCA), it becomes evident that the traditional reliance on cumulative explained variance as a sole metric for model efficacy is insufficient. Surprisingly, high predictive accuracy can be achieved with relatively low cumulative explained variance, highlighting the presence of a minimal subset of “key features” critical for delineating the anti-aging properties of small molecules.

### 4.2 Discovery of Anti-aging Compounds in *C.elegans*

Significantly, our work has led to discovery of some anti-aging compounds: for the first time, we demonstrated the lifespan-extending properties of Thymoquinone, Polyphyllin VI, Medrysone, and Praeruptorin C in the model organism, *C. elegans*. This set of experiments establishes these compounds as potential agents in anti-aging research and underscores the utility of our predictive models.

For those compounds that showed less pronounced effects, it is plausible that the chosen concentrations may have influenced the outcomes. Future studies should consider adjusting the dosages, potentially lowering the concentrations, to refine our understanding of their bioactive properties.

Thymoquinone, historically known for its anti-inflammatory and antioxidant properties and extracted from Nigella sativa or black cumin, now adds lifespan extension in *C. elegans* to its list of beneficial effects. To further validate its role in enhancing stress resistance, we conducted additional heat shock stress experiments at 35°C. Unlike experiments performed at 37°C, the lower temperature of 35°C allowed for a prolonged response period (see Figure S4A).

In addition to thermal stress, we also investigated its capabilities in combating oxidative stress. Thymoquinone was tested against a standard antioxidant, Salidroside, and showed superior efficacy in mitigating oxidative damage. These experiments involved exposing *C. elegans* to environments that induce oxidative stress and measuring the survival and health metrics compared to controls treated with Salidroside (see Figure S4B). The results indicated that Thymoquinone not only extends the lifespan but also enhances the worm’s ability to withstand oxidative environments, surpassing the performance of established antioxidants.

Similarly, Polyphyllin Ⅵ, a saponin derived from the medicinal plant Paris polyphylla and recognized for its anticancer properties, has now been shown to also enhance longevity. Medrysone, typically known for its anti-inflammatory effects in medical treatments, has revealed a new facet of its benefits by demonstrating potential anti-aging effects in *C. elegans*, too. This discovery opens up new possibilities for its therapeutic use and suggests that there are more beneficial properties waiting to be uncovered.

Lastly, Praeruptorin C, originating from the roots of Peucedanum praeruptorum and valued for its cardiovascular benefits, has also shown promise in extending lifespan. This new aspect of its activity enhances its profile as a multifunctional bioactive compound.

### 4.3 Performance and Application of Short Fingerprints

Short fingerprints offer distinct advantages in molecular recognition and drug screening, particularly when optimized with advanced dimensionality reduction techniques such as Principal Component Analysis (PCA) and Kernel PCA (KPCA). Short fingerprints not only overcome the limitations of binary fingerprints in information expression but also provide more refined feature descriptions while retaining enough information to ensure robust predictive accuracy. This capability makes them ideal for rapidly screening huge compound libraries to identify candidates with potential activity.

In this study, short fingerprints processed using PCA and further refined by KPCA not only exhibited outstanding performance but also maintained considerable stability in crucial metrics such as AUC and F1 scores. This stability is crucial in drug discovery since it ensures that these fingerprints perform consistently across diverse chemical spaces and biological activity datasets. Furthermore, short fingerprints efficiently minimize noise in complex data processing, making the model more sensitive to actual biological activity signals.

An essential advantage of using short fingerprints in practical applications, especially when enhanced by KPCA, is their ability to achieve a high F1 score. In drug screening, balancing precision and memory through a high F1 score is as critical as individual accuracy or recall rates, because it ensures that both the presence of false positives and false negatives are minimized. This balanced metric is crucial in the early stages of drug discovery because it ensures that more compounds with genuine biological activity are accurately identified and subsequently validated, optimizing the efficiency and effectiveness of the screening process.

### 4.4 Using the ElixirFP Package to Enhance Molecular Fingerprint Analysis and Interpretation

This article presents ElixirFP, a novel tool for computational drug discovery. ElixirFP, created as a pivotal component of our ongoing research endeavors, emerges as a breakthrough software package poised to revolutionize molecular fingerprint analysis. Its versatile functionality and enhanced interpretability provide researchers with unparalleled insights into chemical properties and interactions.

ElixirFP stands out by offering a comprehensive suite of capabilities, including the generation of three fundamental types of molecular fingerprints: Morgan, Topological, and MACCS. Each fingerprint type provides a distinct perspective on molecular characteristics, allowing researchers to adjust their analysis to the specific requirements of their studies. Moreover, ElixirFP’s advanced algorithms expertly manage the complex balance of information retention and computing efficiency, facilitating the determination of optimal fingerprint lengths for maximal analytical efficacy.

A pivotal feature distinguishing ElixirFP is its provision of importance scores for individual fingerprint bits, which is especially notable in MACCS fingerprints. These scores are excellent guidelines for elucidating the molecular components that exert significant influences on biological activity. ElixirFP significantly enhances result interpretability by providing insights into the essential chemical structures predictive of activity, allowing researchers to make more informed decisions in drug discovery.

The integration of Kernel PCA (KPCA) with the Gaussian Radial Basis Function (RBF) kernel represents a notable advancement within ElixirFP. Using this sophisticated approach, ElixirFP provides weighted fingerprints based on derived importance scores, thereby capturing the intricate nonlinear relationships inherent within the data. This refinement substantially improves both the accuracy and interpretability of the fingerprints, revealing new paths for precise analysis and interpretation in drug discovery research.

In essence, ElixirFP fundamentally transforms drug discovery procedures by offering a flexible range of features designed to tackle the complex issues faced in molecular fingerprint analysis. The heightened interpretability and analytical efficacy of this technology enable researchers to obtain unparalleled insights into the chemical characteristics and interactions, thereby significantly speeding up the process of drug discovery and development.

## Acknowledgments

Special thanks to Prof. Jau-Shyong (John) Hong for his invaluable advice on experimental design and Dr. Shunqi Liu for her assistance with statistical analysis. We also extend our appreciation to the laboratory technicians for their diligent work in maintaining the *C. elegans* cultures and to the administrative staff for their support throughout the project.

## Author Contributions

Conceptualization, B.X. and Y.P.; methodology, Y.P. and H.C.; software, Y.P.; validation, F.Y., W.X., Z.H., J.Z., Y.G, and Y.L.; formal analysis, J.Z., Y.G, and Y.L.; investigation, F.Y., W.X.; resources, B.X., F.Y., J.N., G.S. and J.Y.; data curation, Y.P.; writing—original draft preparation, Y.P. and H.C.; writing—review and editing, F.Y., H.C. A.N.E., J.N. and H.L.; visualization, G.L., G.S.; supervision, J.Y. and B.X.; project administration, B.X. and J.N.; funding acquisition, B.X., J.Y., J.N. and G.L. All authors have read and agreed to the published version of the manuscript.

## Funding

This research was supported in part by the 2021 Research Start-up Fund Fresh Wave (Central Finance Special, Grant No. Y030212059003033 to B.X.), the Leading Principal Investigator of Beijing High-level Public Health Technical Talents Construction Project (Grant No.02-03 to G.L.), the Academic Leader of Beijing High-level Public Health Technical Talents Construction Project (Grant No.02-08 to J.N.), and the Sichuan Science and Technology Program (Grant No. 2022ZYD0076, No. 2023YFS0050, No.2024YFHZ0009 to J.Y.)

## Conflicts of Interest

The authors declare no conflict of interest.

## Ethics approval and consent to participate

Not applicable.

## Availability of data and materials

The datasets analyzed during the current study are available via: https://github.com/Marissapy/ElixirSeeker-ElixirFP

## Notes

https://github.com/Marissapy/ElixirSeeker-ElixirFP

### Competing Interest Statement

The authors have declared no competing interest.

