## Supplementary Figures for "ElixirSeeker: A Machine Learning Framework Utilizing Attention-Driven Fusion of Molecular Fingerprints for the Discovery of Anti-Aging Compounds"

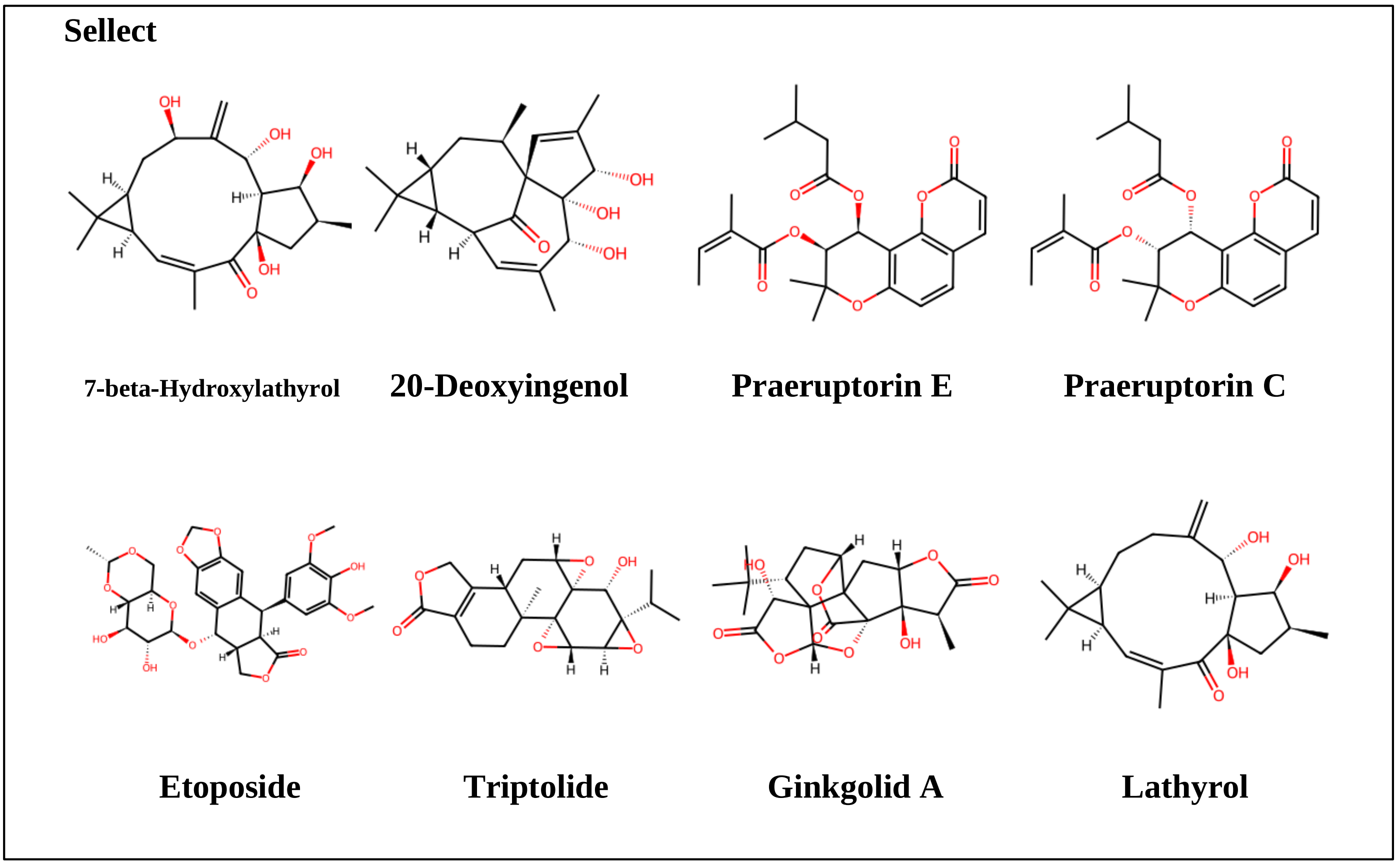


**Fig. S1 TOP 8 moleculars in Sellect compounds database.**

**FDA Approved**

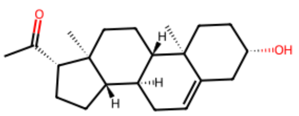

Pregnenolone

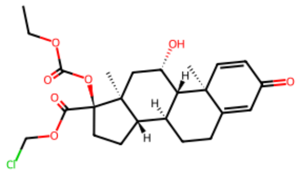

Loteprednol etabonate

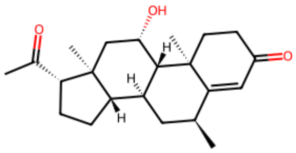

Medrysone

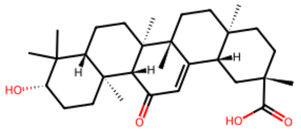

18alpha-Glycyrrhetinic acid

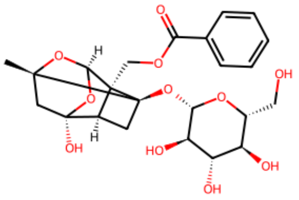

Paeoniflorin

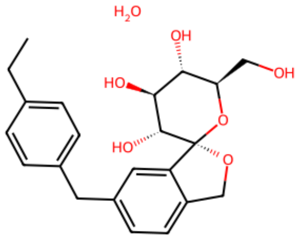

Tofogliflozin hydrate

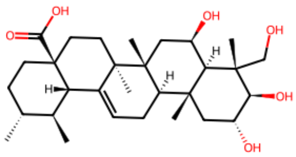

Madecassic acid

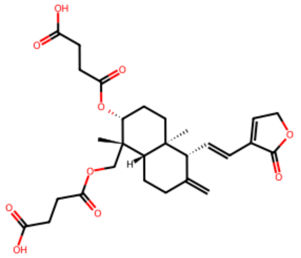

Dehydroandrographolide succinate

**Fig. S2 TOP 8 moleculars in FDA-Approved compounds database.**

**TCM**

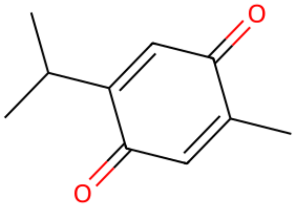

Thymoquinone

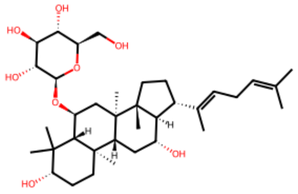

Ginsenoside Rh4

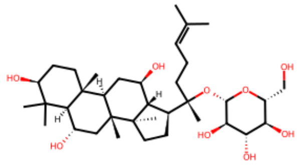

Ginsenoside F1

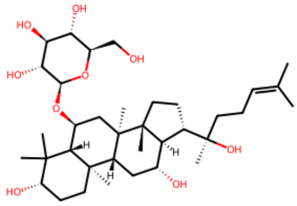

20R-Ginsenoside Rh1

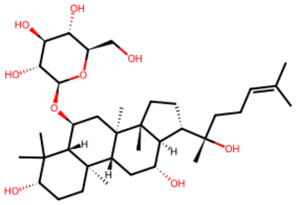

Ginsenoside Rh1

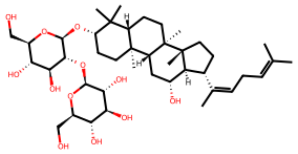

Ginsenoside Rg5

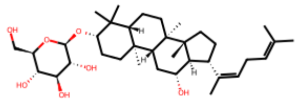

Ginsenoside Rh3

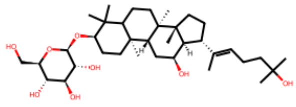

Pseudo-ginsenoside Rh2

**Fig. S3 TOP 8 moleculars in TCM compounds database.**

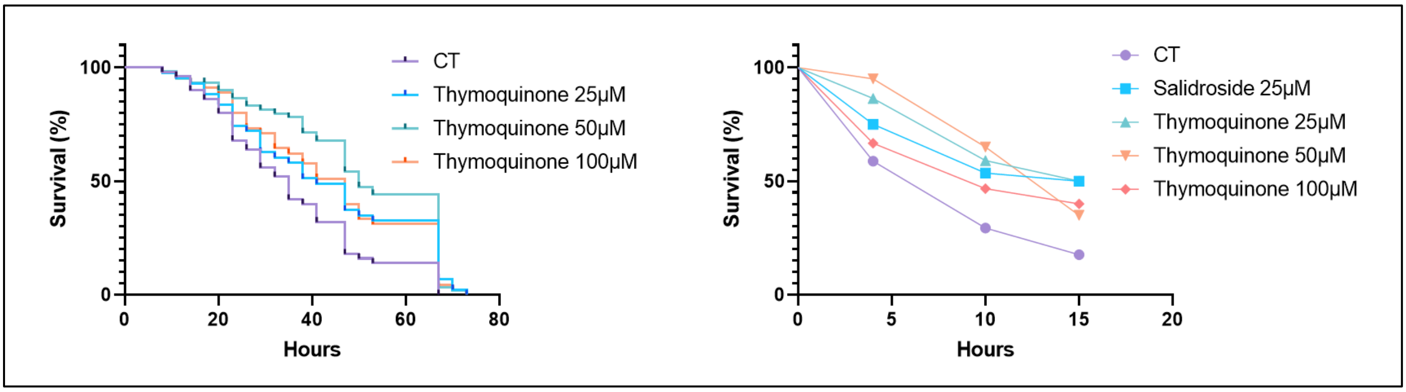


**Fig. S4 Stress Response Assays in C. elegans Treated with Thymoquinone.** A: Survival curves of C. elegans under heat shock conditions at 35°C, comparing control treatment (CT) with varying concentrations of Thymoquinone (25 μM, 50 μM, and 100 μM). B: Oxidative stress assay results showing the survival of C. elegans exposed to oxidative stress over time.
